# BEL Commons: an environment for exploration and analysis of networks encoded in Biological Expression Language

**DOI:** 10.1101/288274

**Authors:** Charles Tapley Hoyt, Daniel Domingo-Fernández, Martin Hofmann-Apitius

## Abstract

The rapid accumulation of knowledge in the field of systems and networks biology during recent years requires complex, but user-friendly and accessible web applications that allow from visualization to complex algorithmic analysis. While several web applications exist with various focuses on creation, revision, curation, storage, integration, collaboration, exploration, visualization, and analysis, many of these services remain disjoint and have yet to be packaged into a cohesive environment.

Here, we present BEL Commons; an integrative knowledge discovery environment for networks encoded in the Biological Expression Language (BEL). Users can upload files in BEL to be parsed, validated, compiled, and stored with fine-granular permissions. After, users can summarize, explore, and optionally shared their networks with the scientific community. We have implemented a query builder wizard to help users find the relevant portions of increasingly large and complex networks and a visualization interface that allows them to explore their resulting networks. Finally, we have included a dedicated analytical service for performing data-driven analysis of knowledge networks to support hypothesis generation.

This web application can be freely accessed at https://bel-commons.scai.fraunhofer.de.

## Introduction

There exists a variety of modeling languages, data formats, and analytical tools for systems and networks biology. Among the most popular modeling languages are the Biological Pathways Exchange [BioPAX; 1], Systems Biology Markup Language [SBML; 2], Systems Biology Graphical Notation [SBGN; 3], and Biological Expression Language [BEL; 4]. BioPAX captures metabolic, signaling, molecular, gene-regulatory, and genetic interaction networks; SBML captures mathematical models of biochemical networks, cellular signaling, and metabolic pathways; SBGN provides a graphical representation of ideas from BioPAX and SBML; and BEL captures qualitative causal and correlative relations between biological entities across multiple scales (e.g., *-omics*, pathway, cellular, phenotypic) with accompanying biological and contextual annotations. While each modeling language has their own domain-specific syntax and semantics, each facilitates assembling biological relations into networks. For example, BEL formalizes relations as triplets each composed of a subject, predicate, and object in order to generate pathways and networks when the object from one relation is again used as the subject of another. We refer to Saqi *et al*. [5] for a more thorough comparison of the applicabilities of various modeling languages.

Currently, these modeling languages and their related analytical tools require deep knowledge of computer programming to use and are generally inaccessible to a wider audience of biologists and clinicians. With the explosion of data and knowledge in the biomedical domain, it is paramount to develop tools that foster collaboration between groups of scientists with different backgrounds and skill sets who are working towards similar goals. Already, there are multiple freely available, web-based tools for systems and networks biology with varying focuses on creation, revision, curation, storage, integration, collaboration, exploration, visualization, and analysis. Below, we provide a brief review of services appropriate for each focus.

Because of the accelerating throughput of scientific publication in the biomedical domain, several general workflows (e.g., REACH [6], TRIPS [7], TEES [8], MedScan [9]) and several BEL-specific workflows (e.g., BELIEF [10], BELMiner [11], BelSmile [12], BELTracker [13]) have been developed to automate biological relation extraction. The limits of the precision and recall of automated techniques, the applicability domains of different modeling languages, and the need for expert input motivated the development of semi-automatic and manual curation interfaces (e.g., SBV Improver [14], BELIEF Dashboard [15], WikiPathways [16]) to drive crowdsourced creation, revision, and curation of knowledge networks.

While many useful knowledge resources (e.g., miRTarBase [17], CTD [18], KEGG [19]) still disseminate their data in non-standard formats, the use of the aforementioned modeling languages has become much more common as in the case of the integration effort of Pathway Commons [20]. Other web tools (e.g., NDEx [21], GraphSpace [22]) provide the ability to upload, store, share, and distribute networks while remaining agnostic to format. Most of these resources also include network visualization, layout, and exploration with PathVisio [23], Cytoscape [24], Cytoscape.js [25], as well as in-browser navigators.

Numerous algorithms and analyses have been published for systems and networks biology, but most are bespoke due the heterogeneous nature of their underlying knowledge networks, data sets, and the scientific questions motivating their development. An exception lies with gene set enrichment analysis: a technique for finding gene sets, pathways, and networks in which a query gene set (e.g., a list of differentially expressed genes) is over- or under-represented [26]. Its simplicity has lead to its implementation and inclusion in several web applications as well as several applications to reveal patterns of dysregulation in *-omics* data sets as exemplified by the Enrichment Map Cytoscape Plugin [27, 28] with Pathway Commons as well as GSEA [29] with MSigDB [30].

Recently, BEL has been successfully used as a semantic and modeling framework for multi-scale and multi-modal knowledge in order to investigate the aetiology of complex neurodegenerative diseases as shown by Domingo-Fernández *et al*. with the NeuroMMSig Mechanism Enrichment Server [31]. While the list of published BEL-specific algorithms is currently short (e.g., Reverse Causal Reasoning [32], Network Perturbation Amplitude [33], etc.), recent developments in the BEL software ecosystem have improved the accessibility and utility of BEL and have motivated its wider adoption [34]. Unfortunately, unlike many of the other focuses of web applications, algorithms have remained confined to use by bioinformatics and inaccessible to a wider audience of researchers across disciplines. Last, but not least, the ecosystem of BEL-specific web applications is small, and does not include a service for parsing, validating, compiling, and converting BEL.

There are still several unmet needs for users that motivate the development of new web applications. Generally, there is still the need to enable complex exploration and visualization as well as to make algorithms and analyses generally accessible and reusable. Specifically to BEL, there is a need to make parsing, validating, compiling, and converting facile and user-friendly. Finally, an integrative knowledge discovery environment that comprises many of the previously mentioned features would be greatly beneficial to the BEL and overarching scientific community. Here, we present BEL Commons, a web application that addresses these unmet needs and is a first attempt at building such an environment.

## Implementation and Components

The user interface of BEL Commons integrates several features from the variety of previously mentioned web applications for systems and networks biology (**Figure 1**). It contains five main components: 1) the network uploader, where users can upload, parse, validate, and compile BEL as well as generate several summaries of its contents; 2) user rights and project management, where users can share their networks with various granularities; 3) the query builder, where users can interactively query networks and make transformations; 4) the biological network explorer, where users can visualize, explore, and further modify networks; and 5) the analytical service, where users can run heat diffusion experiments with *-omics* data. Below, we elaborate on their functionalities and typical use cases. Implementation details can be found in the Supplementary Information.

**Figure 1:**
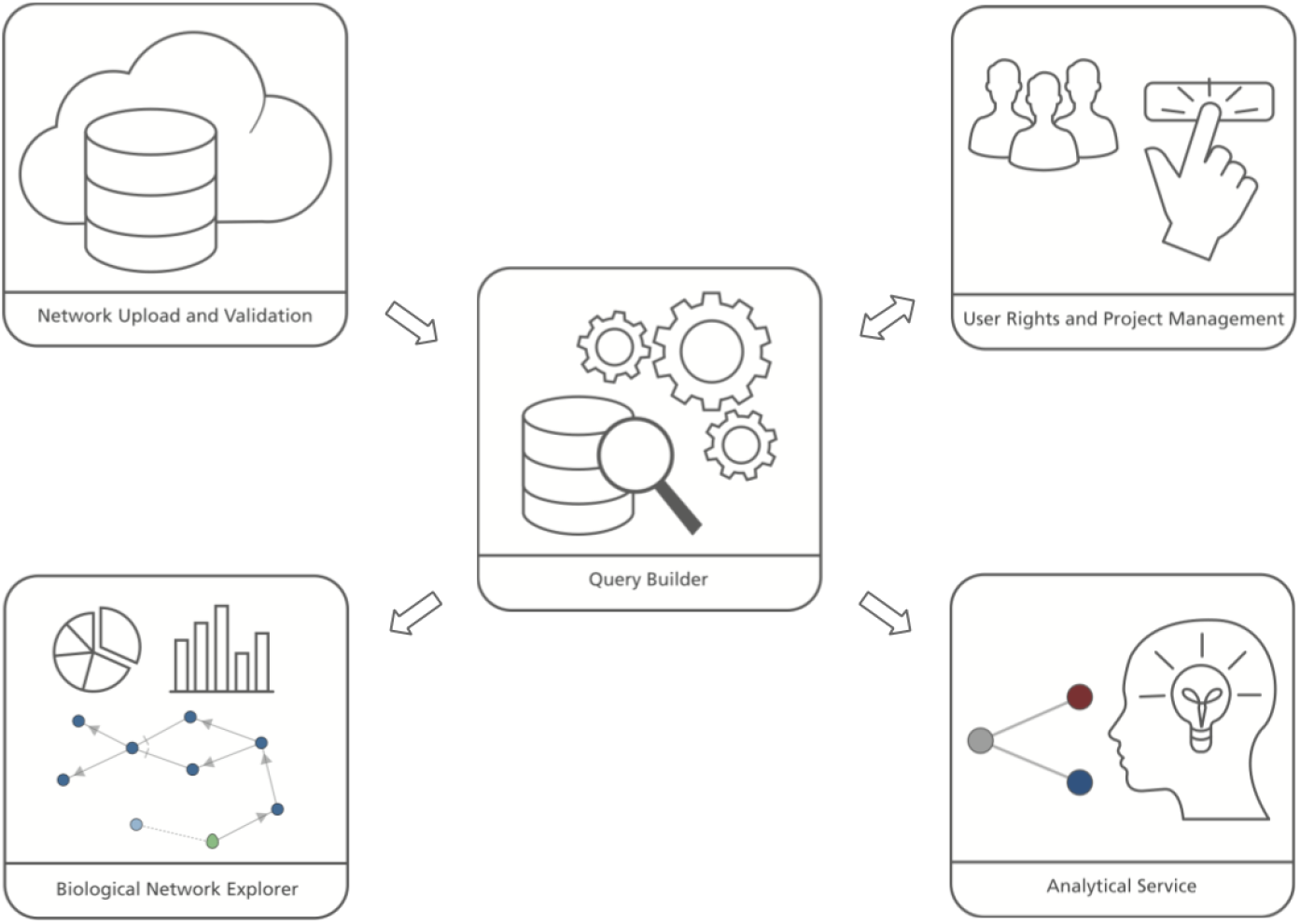
BEL Commons comprises several components: 1) the network uploader and validator, 2) user rights and project management, 3) the query builder, 4) the biological network explorer, and 5) the analytical service.

### Network Uploader

The first point of entry for many users of BEL Commons will be through its BEL uploader, which allows users to choose a file from their computer to upload and to toggle common parsing and compilation parameters. After submitting, users’ files are sent to an asynchronous task queue, implemented with RabbitMQ (https://www.rabbitmq.com) and Celery (http://www.celeryproject.org), that performs parsing, validation, and compilation with PyBEL [34] in the background. Errors and warnings encountered during parsing are enumerated, statistics over the resulting network are produced, biological network motifs are identified, and finally, the submitter is notified upon completion.

The parsing errors and warnings are categorized first as syntactic or semantic then with much more detail, as described in on the BEL Commons help page (https://bel-commons.scai.fraunhofer.de/help/parser). Each is presented with provenance information including the line, line number, and position so curators can quickly make changes. Recurring errors and warnings are identified and grouped separately to allow curators to quickly make impactful improvements. Finally, a faceted search is presented for situations where an overwhelming number of errors and warnings are present.

The statistical summary (**Figure 2A**) of the network presents information about the contents of the network and also network-theoretic measurements of the full network. Several charts are generated depicting the types and number of nodes, edges, modifications, namespaces, annotations, and citations existing in the network. Furthermore, scalar values describing network properties such as network density, average node degree, and node overlap with other networks in BEL Commons using the Szymkiewicz-Simpson coefficient are listed.

**Figure 2:**
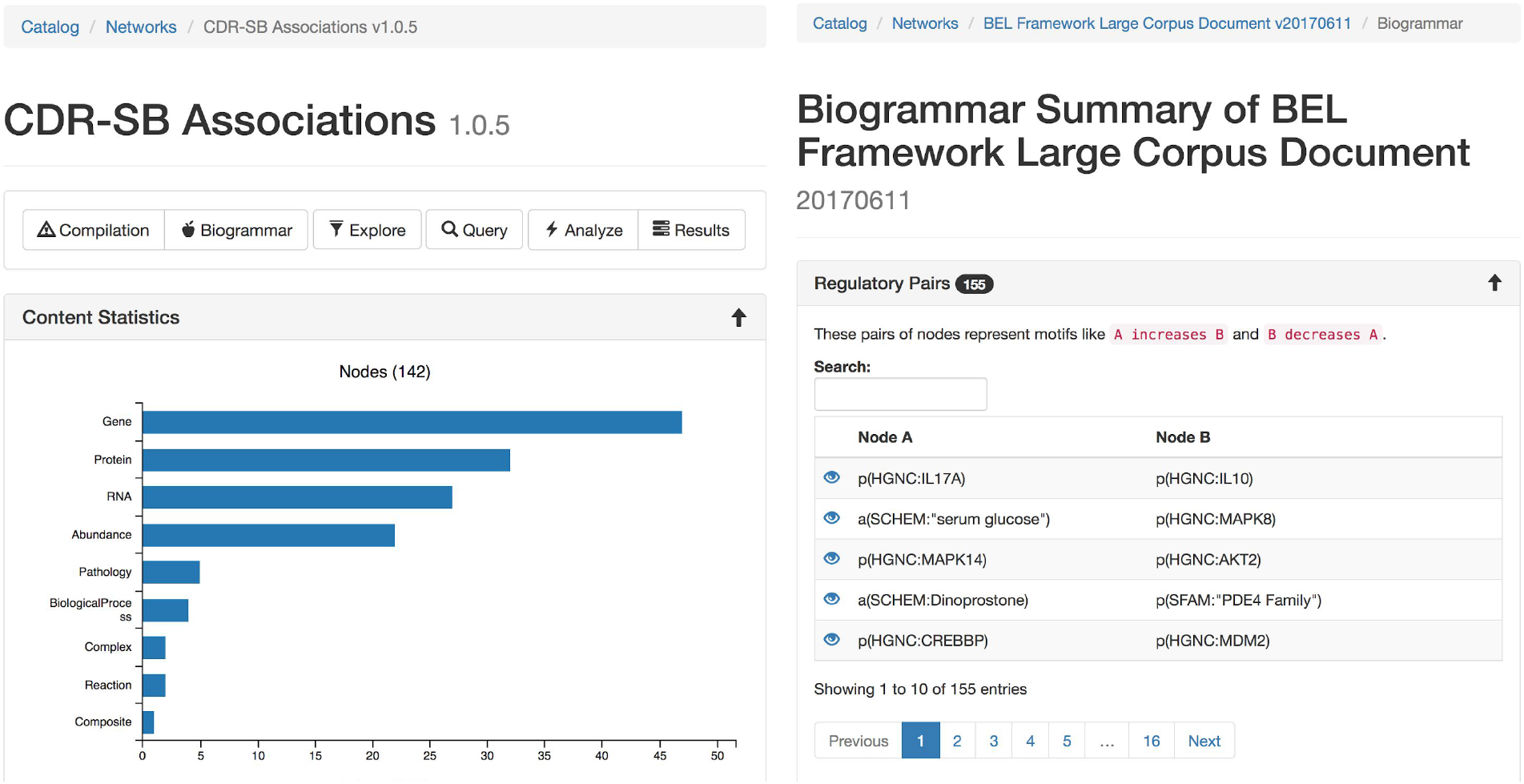
The statistical (A - left) and biogrammar (B - right) summary pages.

The “biogrammar” summary (**Figure 2B**) of the network presents an analysis of network motifs generalized from the analysis of transcriptional network motifs presented by Alon [35] for use with knowledge networks. BEL Commons focuses mainly on simple motifs that are informative to the robustness, correctness, and applicability of a given knowledge work. The most simple motif is the contradictory pair, where knowledge has been curated stating *A increases B* but also *A decreases B*. Searching this motif in NeuroMMSig identified the contradiction that RB1 has been shown to increase the transcriptional activity of E2F4 by Li *et al*. [36] but also that it decreases the transcriptional activity of E2F4 [37]. Another motif is an inconsistent negative correlation triple, where knowledge has been curated stating *A negatively correlates with B, B negatively correlates with C*, and *C negatively correlates with A*. After, several factors can be used to estimate the confidence of the correctness and applicability of the statements such as biological context (e.g., cell line, tissue, disease), reference type (e.g., experimental paper, review paper, database), text location (e.g., abstract, introduction, methods, discussion), or the date of publication. BEL Commons currently only identifies pre-defined, small motifs containing two or three nodes, but could be extended to automatically find larger ones with the caveat that their biological meanings are more difficult to interpret.

### User Rights Managements and Collaboration

Upon BEL upload, users are presented with the option to make the resulting network either public or private. Networks can be uploaded privately for use during research then later released publicly to accompany a publication and share their work with the scientific community. The network catalog (**Figure 3A**) dynamically shows users only networks for which they have the appropriate permissions.

**Figure 3:**
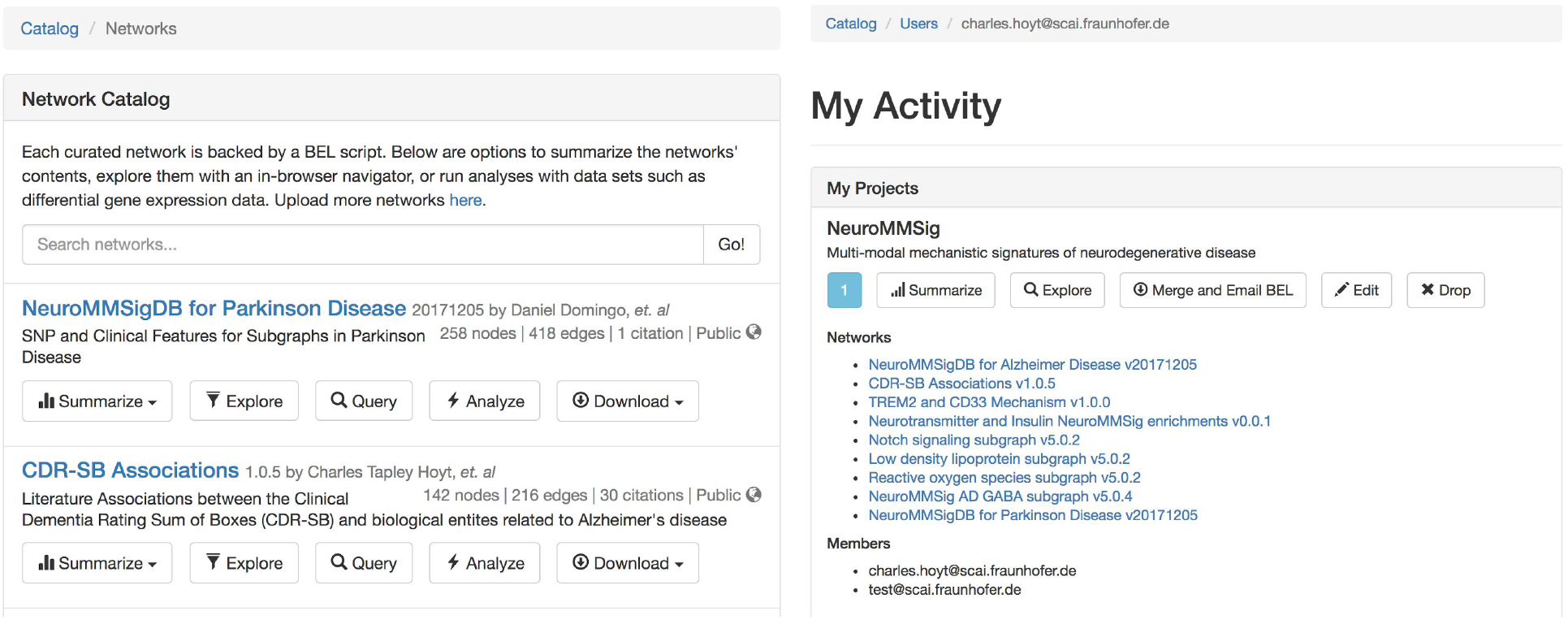
The network catalog (A - left) and user activity page (B - right).

Users can create projects that allow for multiple users to mutually share networks. For example, a curation project in a given disease area could contain networks generated from the efforts of multiple curators. Projects can generate a merged network that can be summarized, explored, analyzed, and exported with the same tools available for stand-alone networks. Users can access their private activity page (**Figure 3B**), which provides a global summary over their projects, networks, queries, data sets, and experiments.

We do not presume all users plan to produce their own BEL, especially with the growing number of both general and context-specific publicly-available resources. In light of this, we have included several of these resources in BEL Commons for these users (**Table 1**). The catalog of networks for which users have the appropriate permissions can be accessed directly from the home page of BEL Commons.

**Table 1.** Statistics over a selection of the resources publicly initially available in BEL Commons. These numbers are accurate to the best of our ability, but may not reflect nominal values from their sources depending on the ability of PyBEL to parse their contents.

| Resource | Networks | Nodes | Edges | Citations |
| --- | --- | --- | --- | --- |
| Selventa Example Corpora (http://resources.openbel.org) | 5 | 16339 | 36971 | 5083 |
| Causal Biological Networks Database [38] | 138 | 5343 | 28766 | 4580 |
| NeuroMMSig [31] | 8 | 1411 | 3221 | 201 |

### Query Builder

While networks arising from BEL can be readily merged, it becomes increasingly difficult to visualize and explore large networks and combinations of networks. The query builder assists users in generating powerful, precise, and expressive queries that find the most relevant and interesting subnetworks in three steps: 1) users search and select relevant networks, 2) users generate a subnetwork specifying the most interesting nodes, edge annotations, and references with their preferred *seeding method(s*), and 3) users select transformations (e.g., enrichments, selections, filters) to apply to the resulting subnetwork.

The first step in the query builder allows users to select networks relevant to their scientific questions. While it still remains computationally feasible to query over a merged view of the entire catalog of networks, users have the opportunity to pre-select the most relevant networks on the basis of their specificities towards target diseases areas, curation methods, or any other appropriate criteria described in their metadata. Additionally, other high-quality structured knowledge resources such as protein families [39, 40], biochemical reactions [41, 42], gene orthologies (e.g., Entrez Gene [43], MGI [44], RGD [45], etc.) can be included to enrich networks from curated BEL. Later, we show how this novel feature can be used to enrich networks as a pre-processing step to connect disparate components before analysis.

The second step in the query builder allows users to generate subnetworks based on nodes, edge annotations, and references of interest by using one or several “seeding methods.” An example seeding method that most network-related web applications implement is the retrieval of the N^th^ neighbors of a given node or set of nodes. Because neighborhood queries are often insufficient to capture complex biology, BEL Commons implements several additional seeding methods, enumerated in **Table 2**, that allow users to take advantage of the directionality, polarity, and rich annotations inherent to networks from BEL. Using these seeding methods, the query builder allows scientists to ask scientific questions like the one proposed in the following scenario; the leukemia drug, nilotinib, triggers cells to remove faulty components, including ones associated with several brain diseases [46]. In 2015, the Georgetown University Medical Center published findings that the drug had a therapeutic effect on patients in Alzheimer’s and Parkinson’s diseases [47]. Though the drug’s mechanism of action is currently unknown, a path search between nilotinib and these diseases suggests it could be by decreasing phosphorylation of the Tau protein, which may have a therapeutic effect in both disease contexts, through inhibition of ABL1 [48, 49].

**Table 2.** Seed methods available in the query builder.

| Seed Method | Description |
| --- | --- |
| $N^{\text{th}}$ Neighbors | Induces a subnetwork over nodes in paths of length less than or equal to N from any query node, terminating at any node, including ones not included in the query |
| Upstream Subnetwork | Induces a subnetwork over nodes with causal edges targeting the query nodes then repeats a second time for that subnetwork in order to include a second layer. Finally, induces all causal edges between resulting nodes |
| Downstream Subnetwork | Induces a subnetwork over nodes with causal edges originating from the query nodes then repeats a second time for that subnetwork in order to include a second layer. Finally, induces all causal edges between resulting nodes |
| Shortest Paths | Induces a subnetwork over all nodes in the shortest paths between any pair of query nodes, implemented by NetworkX [50] |
| All Paths | Induces a subnetwork over all nodes in all of the paths (length less than seven) between any pair of query nodes, implemented by NetworkX [50] |
| Provenance | Builds a subnetwork from all edges with provenance from articles with the given PubMed identifiers |
| Authors | Builds a subnetwork from all edges from articles written by authors with the given names |
| Annotations | Builds a subnetwork from all edges matching given biological or contextual annotations |

The third step in the query builder allows users to specify of transformations (e.g., enrichments, selections, filters) to apply to network resulting from the assembly in the first step then the seeding in the second step. Users may select basic transformations, such as deleting single nodes or edges, to more complex transformations such as selecting a subnetwork only consisting of causal edges. BEL Commons dynamically loads transformation functions from PyBEL such that new functions, pipelines, and workflows for processing networks can be written quickly and made available to users. A full list is available at https://bel-commons.scai.frannhofer.de/help/query-builder.

Each query is saved with a unique identifier such that queries can be rerun, shared, merged, and compared. Rather than storing the results of queries, the selection, seeding, and transformations are stored as a “transaction” so that they can be applied to new assemblies, for example, when a network is updated. Effectively, queries correspond to an experimental protocol for processing raw networks before visualization, exploration, analysis, and interpretation. However, the construction of a query is not the end of its life. The next section summarizing the biological network explorer describes how queries can be extended and evolve during the process of scientific inquiry.

### Biological Network Explorer

The biological network explorer provides users with easy ways to visualize BEL networks, to interpret their underlying structures, to investigate the metadata on nodes and edges, and to interactively update networks as they explore (**Figure 4**). It is built with D3.js (https://d3js.org) to render networks with a force-directed layout algorithm that can be panned and zoomed. Because the complexity of biological networks often limits the utility of automated layouts [51], users can also manually drag and reposition nodes. Furthermore, users can adjust the edge length parameter of the algorithm to rarefy densely grouped nodes and improve readability. The networks are styled with minimum visual clutter and make use of easily-distinguishable colors rather than obtrusive shapes for nodes as well as patterns and colors for different types of edges.

**Figure 4:**
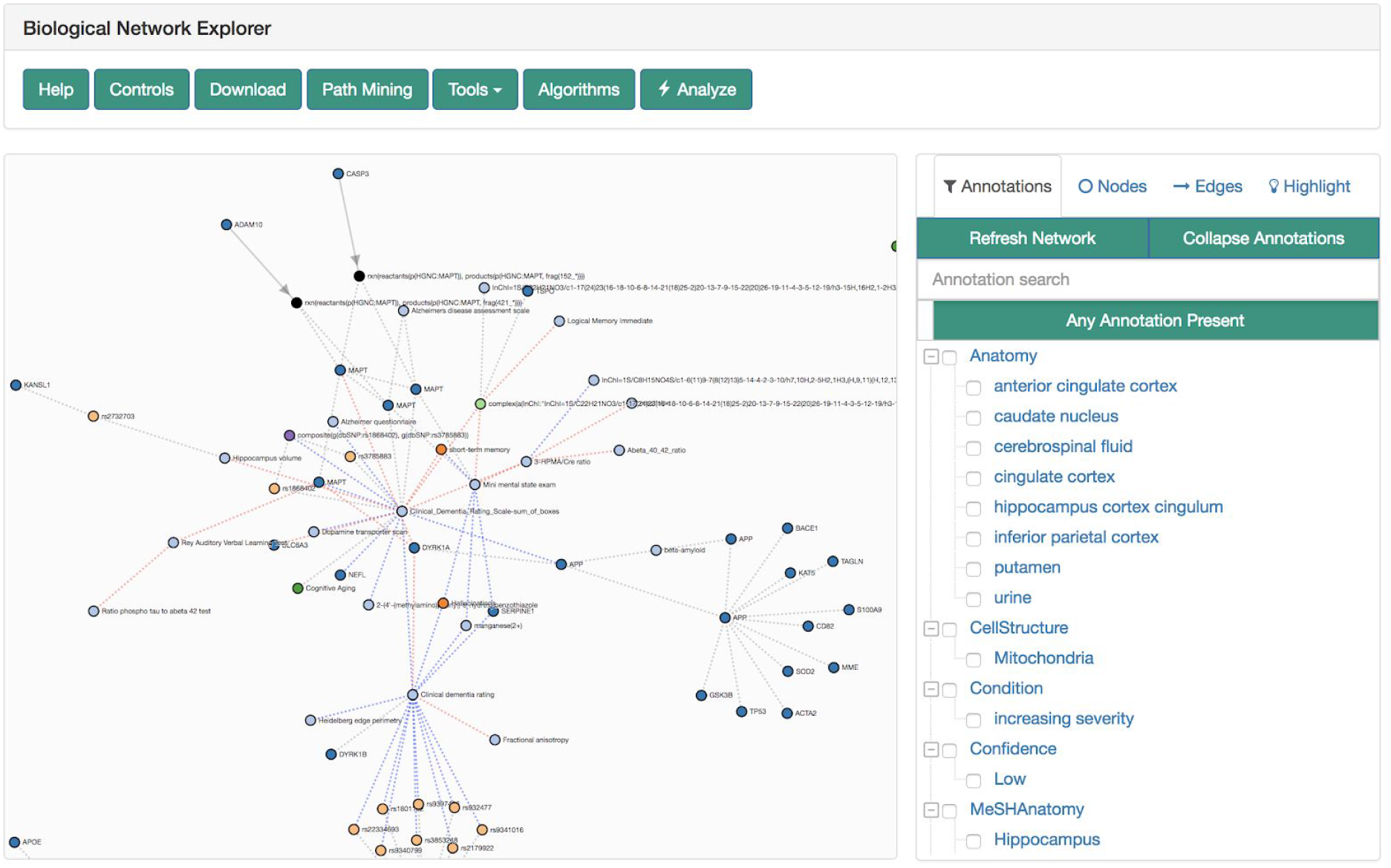
The biological network explorer and related navigation components. Truncated from this image are the node information and query information boxes.

The explorer has several contextual actions registered to the nodes and edges. Users can left-click a node to populate the information box located below the explorer with information from external data sources (e.g., Entrez Gene, ChEBI [52], ExPASy [40], Gene Ontology [53] gathered from Bio2BEL services (https://github.com/bio2bel) and the EMBL Ontology Lookup Service [54]. Alternatively, users can right-click a node to open a contextual menu that enables further modification to the network (e.g., delete the node, add the neighbors of the node to the network) that are interactively appended to the original query used to render the visualization. The contents of the network can also be further modified by the inline query builder, which allows additional transformations to be applied interactively. The query history is displayed at the bottom of the explorer, new changes are highlighted in red, and because queries are stored as transactions, changes can be reverted with an “undo” button.

When an edge is clicked, the information box is populated with relevant citations, evidences, and annotations. Each edge is linked to a voting and commenting system so domain-specific experts, curators, and bioinformaticians can engage in discussion on the correctness and robustness of the chosen representation of knowledge.

To the right of the explorer is the filter tool box, which incorporates a novel approach to filtering and exploring networks using a linked hierarchical explorer of the terminologies/ontologies annotated to the edges in the currently displayed networks. Users can search and select groups of annotations to filter the network. For example, this could be useful to exclude edges asserted from research on cell lines that are not relevant. The filter tool box has three additional tabs; nodes, edges and highlight. Users can either search for specific nodes and edges in their corresponding tabs or use the highlight tab to select nodes and edges with specific properties to highlight in the network.

Above the explorer is the general tool box that includes several additional interactions for exploration, analysis, and export of the network using the serializers described by Hoyt, *et al*. [34]. Notably, it contains a path mining tool that enables path searches between given nodes with fine-granular, configurable settings (e.g., directedness, path search algorithm, application of filters for pathologies, etc.). Hence, it can immediately be used to identify the causal root affecting two nodes, or generate hypothetical links across modes and scales.

The visualization can be further modified by resizing the nodes corresponding to the results of topological or data-driven analyses, such as their degree, betweenness centrality, or by the results of an experiment (e.g. heat diffusion with *-omics* data; see next section) in order to identify novel biological entities.

Finally, there are several alternatives to exploring networks that are too large to render in-browser due to the limits of Javascript-based graphics. First, the network catalog opts to present users with a random subsampling of large networks. If a network in the explorer becomes too big, then the explorer prompts the usage of the filter tool box to identify a more relevant, smaller network. Otherwise, users can export the current network to multiple formats for use in desktop visualization applications.

### Analytical Service

Khatri *et al*. categorized algorithms for analyzing pathways and networks in three types: over-representation analysis, functional class scoring, and pathway topology [26]. Algorithms of each type have been developed for a wide variety of applications, data formats, and network types, but most are difficult to use and few are specific to networks from BEL. The BEL Commons analytical service begins to address this issue by coupling a heat diffusion workflow to the query builder and explorer to create a more seamless user experience.

In the context of network science, heat diffusion refers to annotating a scalar value to each node (i.e., heat) and simulating how it spreads through nodes’ adjacent edges to their neighbors over several iterations. It has been used successfully in systems and networks biology to assess the connectivity of nodes and identify important subnetworks as demonstrated by Leiserson *et al*. with the HotNet2 algorithm [55].

BEL Commons exposes a similar workflow, previously presented by Hoyt *et al*. [34], that has the added behavior based on the polarity of causal edges - when heat crosses a *decreases* edge, its sign is flipped to better capture the aggregate effect of heat flowing from several edges with mixed polarity to a single node. Because BEL contains several entity types, users are presented with the final heats on biological process nodes to assist in interpreting which processes are dysregulated in the experiment. A more detailed description of this method can be found at https://bel-commons.scai.fraunhofer.de/help/heat-diffusion.

Users can upload pre-processed, high throughput *-omics* experiments (e.g., differential gene expression data) and map them to a network either by starting in the network catalog, or through the biological network explorer’s toolbox. The network and *-omics* data are then sent to the task queue to perform the heat diffusion workflow. Upon completion, users are notified via email with a link to the results page that shows statistics and data visualization. Finally, users are also able to overlay the results on the original network in the biological network explorer.

In the following section, the query builder, biological network explorer, and analytical service are used to assess several differential gene expression experiments representing Alzheimer’s disease patients at different disease states using NeuroMMSig networks.

### Application Scenario

This section describes a use case in which a disease-specific network for Alzheimer’s disease is assembled and pre-processed. First, the network is explored with the biological network explorer, second, it is enriched, and finally, it is analyzed with differential gene expression data in order to identify patterns of dysregulation of biological processes that are specific to disease progression stages.

We uploaded several BEL documents describing Alzheimer’s disease pathophysiology generated during the AETIONOMY project (https://www.aetionomy.eu) that were originally stored in NeuroMMSig (https://neurommsig.scai.fraunhofer.de) to BEL Commons. We began by using the query builder to search for and select all of these networks. Next, we used the “Annotations” seeding method (described in **Table 2**) to generate a subnetwork composed of edges that had been annotated in BEL with membership in the following candidate mechanisms of Alzheimer’s disease pathophysiology from NeuroMMSig. We chose the Low Density Lipoprotein Subgraph, the GABA Subgraph, the Notch Signaling Subgraph, and the Reactive Oxygen Species Subgraph because they are dysregulated at different disease stages. Later, we will capture these different progression patterns by running the heat diffusion workflow with stage-specific differential gene expression data.

We used the query builder to apply several transformations to pre-process the network, including: 1) enrichment of the network with the members of all protein families and protein complexes, 2) deletion of nodes having the MGI and RGD namespaces which respectively correspond to genes from mice and rats, 3) removing pathology nodes, which often are uninformative hubs in disease-specific networks, and 4) extracting only causal edges. Finally, we submitted the query and visualized the network.

The biological network explorer showed that there were several disconnected components in the resulting network. Further, there were several cases where the gene, RNA, or protein, such as the GABRA4 gene and RNA, were in different components. We used the tool box above the explorer to apply an additional filter, “Enrich Protein And RNA Origins,” which added the corresponding RNA for each protein then the corresponding gene for each RNA/miRNA in the network. Finally, we applied “Collapse Variants” and “Collapse to Genes” to simplify the network by collapsing all corresponding genes, RNAs, proteins, and their variants to a single node for use with the heat diffusion workflow.

After clicking the “Analyze” button above the explorer, we uploaded three differential gene expression analyses corresponding to Alzheimer’s disease patients in three disease stages (i.e., early, moderate, and severe) from Blalock, *et al*. [GSE28146; 56] pre-processed with GEO2R [57]. We applied the heat diffusion workflow to the previously generated network using each of the three differential gene expression analysis in parallel and displayed the results (i.e., the final heat on each biological process in the network) together with the parallel coordinate display in BEL Commons (**Figure 5**). Because each biological process has a final heat corresponding to experiments for the three disease stages, the parallel plot directly allows interpretation of progression patterns. We used BEL Commons to apply a K-Means clustering with *K*=5 to assist in identifying clusters of biological processes with similar progression patterns, and color them accordingly. In Figure 5, biological processes in *group 0* (blue) tended to decrease only at severe onset of disease (e.g., glial cell differentiation). Conversely, biological processes in *group 1* (orange) tended to increase throughout progression of disease (e.g., Notch signaling pathway). Finally, *group 2* (green) processes remained relatively unchanged (e.g. lipid metabolic process) through the progression of disease and *group 4* (purple) remained consistently elevated (e.g., apoptotic process). The complete results of this experiment as well as a tutorial for reproduction can be found in the supplementary data.

**Figure 5:**
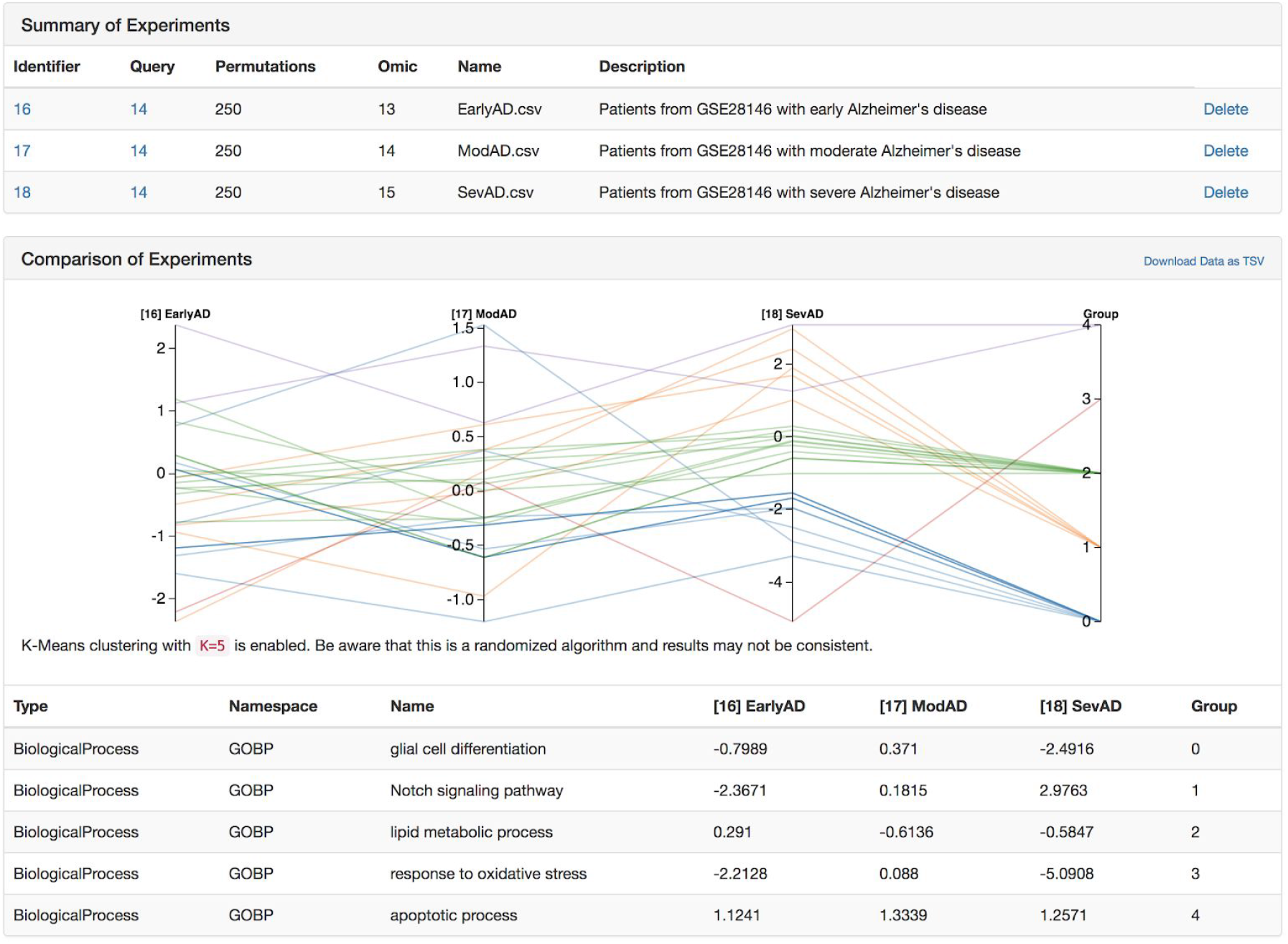
A parallel coordinate plot of the final heats from biological processes from several Alzheimer’s disease-specific networks after running the heat diffusion workflow with differential gene expression data from Blalock *et al*. [56] comparing patients at three stages of Alzheimer’s disease.

While BEL has inherent limits in its temporal expressivity, using temporal data in analysis is an initial attempt to overcome these limits. Complex diseases like Alzheimer’s must be studied with respect to its progression over time, and we believe that workflows like the one presented above could begin to provide insight to the genesis and progression of the disease in order to support patient stratification and precision medicine.

## Discussion

No web application, however feature-rich, will ever satiate the desire and creativity of researchers to generate novel solutions to scientific problems. Even though BEL Commons has taken inspiration from many well-constructed services to build a knowledge discovery environment that enables researchers to explore knowledge and data in new ways, it still shares this limit. However, we are not discouraged, and we hope to make several improvements to BEL Commons in the future.

We would like to improve the interoperability of BEL Commons and the platform build on BEL itself by integrating open authentication systems like ORCID (https://orcid.org) in order to harmonize identification of users across multiple web services and provide reliable provenance for networks, queries, and analyses. We would also like to integrate further tools for converting BEL to RDF in order to connect BEL with other linked data. Further, we would like to improve exporters to other services, notably, NDEx, which have brighter outlooks on sharing and feedback systems. Recent developments in integrating INDRA [58] with PyBEL enables conversion from BioPAX documents to BEL. A future update to BEL Commons will include an option to upload these documents as well.

We would like to integrate BEL Commons with other BEL-specific systems developed with different underlying technologies. First, integrating the BELIEF Dashboard to use the underlying network and edge store from PyBEL would enable a more thorough feedback and curation interface so users could not only vote on the correctness of edges, but fix them directly. Second, the NeuroMMSig Mechanism Enrichment Server will be re-implemented completely with reusable PyBEL code and BEL Commons components in order to advance its goals of achieving patient-stratification by using common algorithms and tools.

## Conclusion

Along with recent improvements in generation of BEL content through text mining (INDRA) and serialization of resources (Bio2BEL project; https://github.com/bio2bel), we believe that BEL Commons will make BEL more accessible to both academic and industrial users. We have made this application freely available at https://bel-commons.scai.fraunhofer.de.

## Authors’ Contributions

C.T.H. and D.D.F. conceived the web application, implemented it, and wrote this manuscript. M.H.A. reviewed the content.

## Acknowledgements

We would like to thank all colleagues that assist in testing and provide feedback in order to improve this work, especially to André Gemünd for his technical assistance. We would also like to thank Scott Colby for making our logos.

